# Decoding neural signals and discovering their representations with a compact and interpretable convolutional neural network

**DOI:** 10.1101/2020.06.02.129114

**Authors:** Arthur Petrosuan, Mikhail Lebedev, Alexei Ossadtchi

## Abstract

Brain-computer interfaces (BCIs) decode information from neural activity and send it to external devices. In recent years, we have seen an emergence of new algorithms for BCI decoding including those based on the deep-learning principles. Here we describe a compact convolutional network-based architecture for adaptive decoding of electrocorticographic (ECoG) data into finger kinematics. We also propose a theoretically justified approach to interpreting the spatial and temporal weights in the architectures that combine adaptation in both space and time, such as the one described here. In these architectures the weights are optimized not only to align with the target sources but also to tune away from the interfering ones, in both the spatial and the frequency domains. The obtained spatial and frequency patterns characterizing the neuronal populations pivotal to the specific decoding task can then be interpreted by fitting appropriate spatial and dynamical models.

We first tested our solution using realistic Monte-Carlo simulations. Then, when applied to the ECoG data from Berlin BCI IV competition dataset, our architecture performed comparably to the competition winners without requiring explicit feature engineering. Moreover, using the proposed approach to the network weights interpretation we could unravel the spatial and the spectral patterns of the neuronal processes underlying the successful decoding of finger kinematics from another ECoG dataset with known sensor positions.

As such, the proposed solution offers a good decoder and a tool for investigating neural mechanisms of motor control.

## 1 Introduction

Brain-computer interfaces (BCIs) link the nervous system to external devices [4] or even other brains [18]. While there exist many applications of BCIs [1], clinically relevant BCIs have received most attention that aid in rehabilitation of patients with sensory, motor, and cognitive disabilities [13]. Clinical uses of BCIs range from assistive devices to neural prostheses that restore functions abolished by neural trauma or disease [2].

BCIs can deal with a variety of neural signals [17, 9] such as, for example, electroencephalographic (EEG) potentials sampled with electrodes placed on the surface of the head [12], or neural activity recorded invasively with the electrodes implanted into the cortex [6] or placed onto the cortical surface [22]. The latter method, which we consider here, is called electrocorticography (ECoG). Accurate decoding of neural signals is key to building efficient BCIs.

Typical BCI signal processing comprises several steps, including signal conditioning, feature extraction, and decoding. In the modern machine-learning algorithms, parameters of the feature extraction and decoding pipelines are jointly optimized within computational architectures called Deep Neural Networks (DNN) [10]. DNNs derive features automatically when trained to execute regression or classification tasks. While it is often difficult to interpret the computations performed by a DNN, such interpretations are essential to gain understanding of the properties of brain activity contributing to decoding, and to ensure that artifacts or accompanying confounds do not affect the decoding results. DNNs can also be used for knowledge discovery. In particular, interpretation of features computed by the first several layers of a DNN could shed light on the neurophysiological mechanisms underlying the behavior being studied. Ideally, by examining DNN weights, one should be able to match the algorithm’s operation to the functions and properties of the neural circuitry to which the BCI decoder connects. Such physiologically tractable DNN architectures are likely to facilitate the development of efficient and versatile BCIs.

Several useful and compact architectures have been developed for processing EEG and ECoG data. The operation of some blocks of these architectures can be straightforwardly interpreted. Thus, EEGNet [8] contains explicitly delineated spatial and temporal convolutional blocks. This architecture yields high decoding accuracy with a minimal number of parameters. However, due to the cross-filter-map connectivity between any two layers, a straightforward interpretation of the weights is difficult. Some insight regarding the decision rule can be gained using DeepLIFT technique [25] combined with the analysis of the hidden unit activation patterns. Schirrmeister et al. describe two architectures: DeepConvNet and its compact version ShallowConvNet. The latter architecture consists of just two convolutional layers that perform temporal and spatial filtering, respectively [23]. Recent study of Zubarev et al. [28] reported two compact neural network architectures, LF-CNN and VAR-CNN, that outperformed the other decoders of MEG data, including linear models and more complex neural networks such as ShallowFBCSP-CNN, EEGNet-8 and VGG19. LF-CNN and VAR-CNN contain only a single non-linearity, which distinguishes them from most other DNNs. This feature makes the weights of such architectures readily interpretable with the well-established approaches [7, 5]. This methodology, however, has to be applied taking into account the peculiarities of each specific network architecture.

Here we introduce another simple architecture, developed independently but conceptually similar to those listed above, and use it as a testbed to refine the recipes for the interpretation of the weights in the family of architectures characterized by separated adaptive spatial and temporal processing stages. We thus emphasize that when interpreting the weights in such architectures we have to keep in mind that these architectures tune their weights not only to adapt to the target neuronal population(s) but also to minimize the distraction from the interfering sources in both spatial and frequency domains.

## 2 Methods

Figure 1 illustrates a hypothetical relationship between motor behavior (hand movements), brain activity, and ECoG recordings. The activity, **e**(*t*), of a set of neuronal populations, *G*_1_−*G*_*I*_, engaged in motor control, is converted into a movement trajectory, *z*(*t*), through a non-linear transform *H*: *z*(*t*) = *H*(**e**(*t*)). The activity of populations *A*_1_× *A*_*J*_ is unrelated to the movement. The recordings of **e**(*t*) with a set of sensors are represented by a *K*-dimensional vector of sensor signals, **x**(*t*). This vector can be modeled as a linear mixture of signals resulting from application of the forward-model matrices **G** and **A** to the task-related sources, **s**(*t*), and task-unrelated sources, **f** (*t*), respectively:

**Figure 1:**
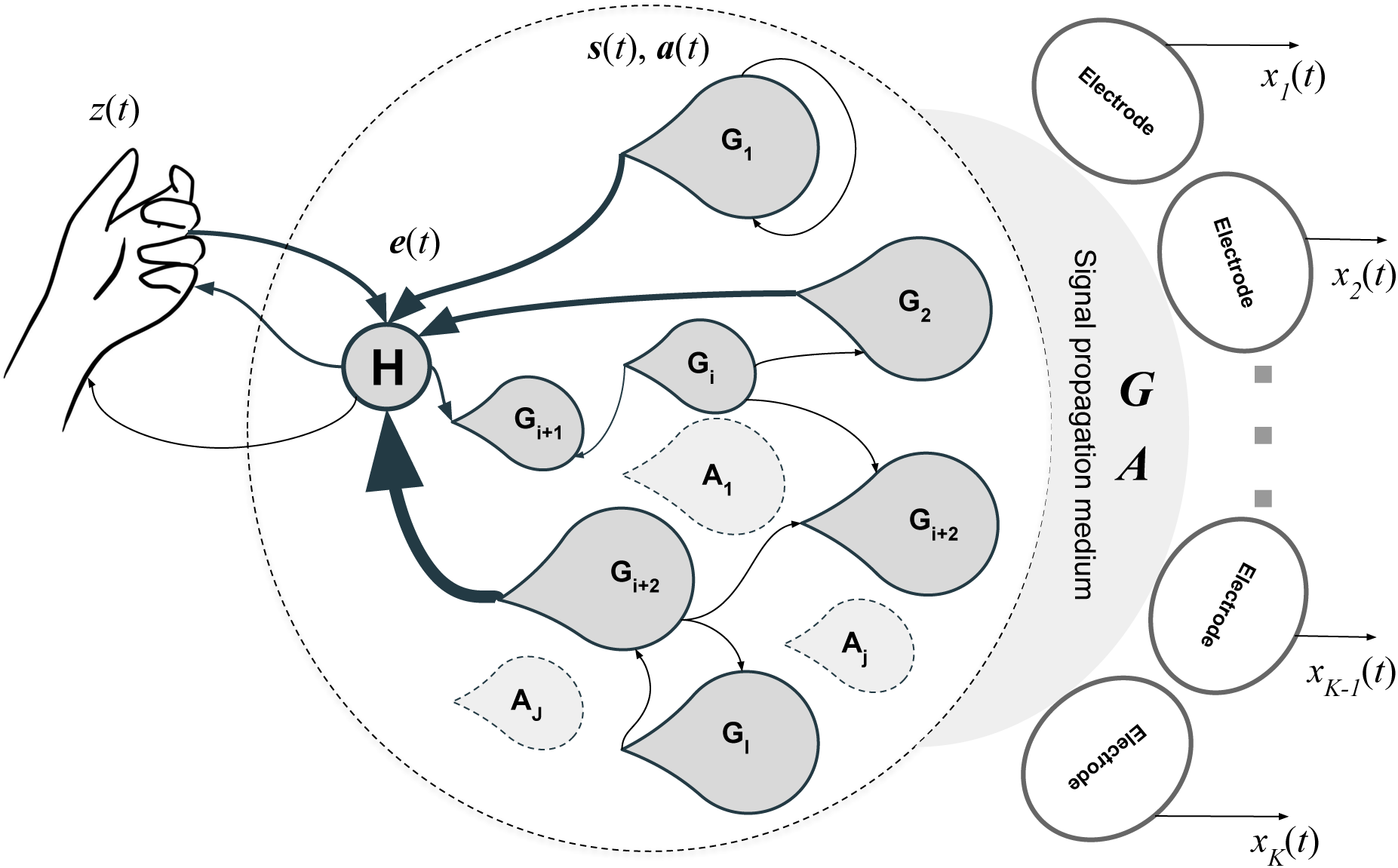
Phenomenological diagram

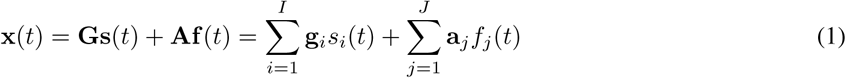

We will refer to the noisy, task-unrelated component of the recording as 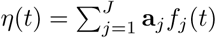.

Given the linear generative model of electrophysiological data, the inverse mapping used to derive the activity of sources from the sensor signals is also commonly sought in the linear form: ŝ(*t*) = **W^*T*^X**(*t*), where columns of **W** form a spatial filter that counteracts the volume conduction effect and decreases the contribution from the noisy, task-unrelated sources.

Neuronal correlates of motor planing and execution have been extensively studied [27]. In the cortical-rhythm domain, alpha and beta components of the sensorimotor rhythm desynchronize just prior to the execution of a movement and rebound with a significant overshoot upon the completion of a motor act [14]. The magnitude of these modulations correlates with the person’s ability to control a motor-imagery BCI [20]. Additionally, the incidence rate of of beta bursts in the primary somatosensory cortex is inversely correlated with the ability to detect tactile stimuli [24] and also affects other motor functions. Intracranial recordings, such as ECoG, allow reliable measurement of the faster gamma band activity, which is temporally and spatially specific to movement patterns [26] and is thought to accompany movement control and execution. Overall, based on the very solid body of research, rhythmic components of brain sources, **s**(*t*), appear to be useful for BCI implementations.Given the linearity of the generative model (1), these rhythmic signals reflecting the activity of specific neuronal populations can be computed as linear combinations of narrow-band filtered sensor data **x**(*t*).

The most straightforward approach for extracting the kinematics, *z*(*t*), from brain recordings, **x**(*t*), is to use con-currently recorded data and directly learn the mapping *z*(*t*) =ℱ(**x**(*t*)). To practically implement it, one needs to parametrically describe this mapping. Here we used a specific network architecture for this purpose. The architecture was constructed in a close correspondence with the observation equation (1) and the neurophysiological description of the observed phenomena illustrated in Figure 1, which facilitated our ability to interpret the results.

### 2.1 Network architecture

The compact and adaptable architecture that we used here is shown in Figure 2. An adaptive envelope extractor is the key component of this architecture. The envelope extractor, a module widely used in signal processing systems, can be implemented using modern DNN primitives, namely a pair of convolutional operations that perform band-pass and low-pass filtering and one non-linearity ReLu(−1) that corresponds to computing the absolute value of the output of the first 1-D convolutional layer. To make the decision rule of this structure tractable, we used non-trainable batch normalization when streaming the data though the structure.

**Figure 2:**
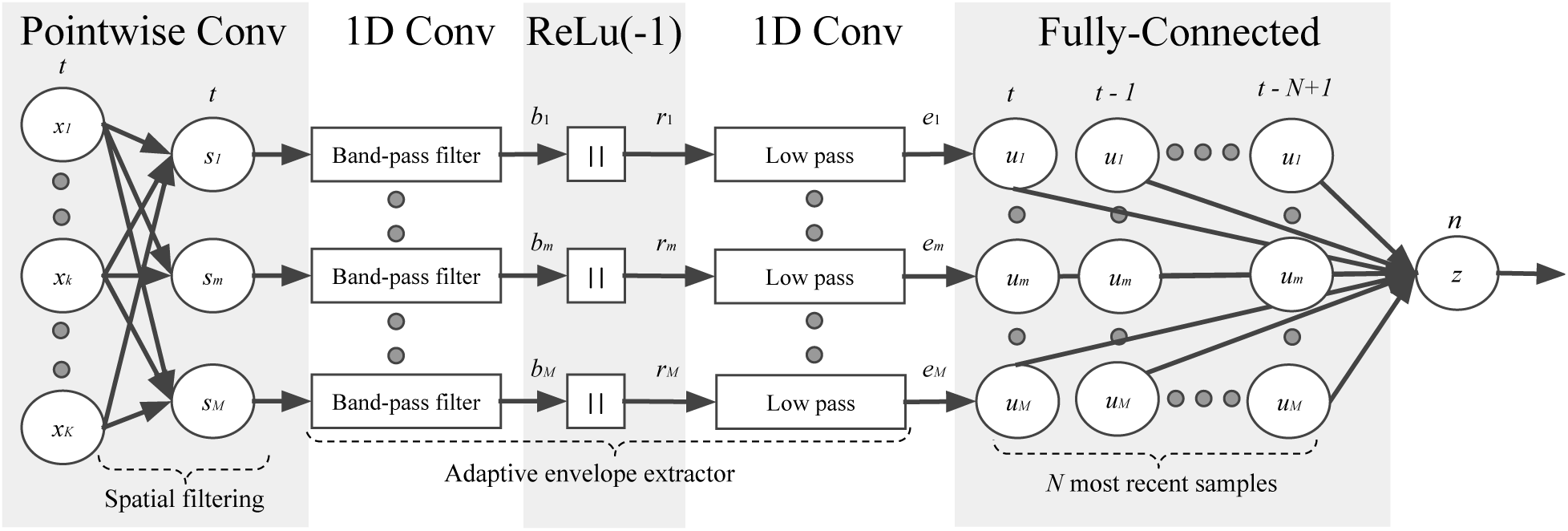
The architecture based on the compact CNN comprises several branches - adaptive envelope detectors, receiving spatially unmixed input signals and outputting the envelopes whose *N* most recent values are combined in the decoded variable *z* by the fully connected layer

In our architecture, the envelope detectors received as an input spatially filtered sensor signals **s**_*m*_ that were calculated by the point-wise convolutional layer. This layer is designed to counteract the volume-conduction processes represented by the forward-model matrices **G** and **A** in our phenomenological model (Figure 1). Next, we approximated operator *H* as a function of the lagged instantaneous power (envelope) of the narrow-band source timeseries. This was done with a fully connected layer that mixed the samples of envelopes, *e*_*m*_(*n*), into a single estimate of the kinematic parameter *z*(*n*).

### 2.2 Two regression problems and DNN weights interpretation

The described architecture processes data in chunks of a prespecified length of *N* samples. We will first assume that the chunk length is equal to the filter length in the 1-D convolution layers. Then, the processing of a chunk of input data, **X**(*t*) = [**x**(*t*), **x**(*t* 1), … **x**(*t* −*N* + 1)], by the first two layers performing spatial and temporal filtering can be described for the *m*-th branch as

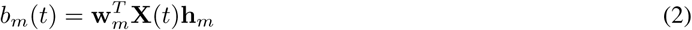

The non-linearity, *ReLu*(− 1), in combination with the low-pass filtering performed by the second convolutional layer, extracts the envelopes of rhythmic signals.

The analytic signal is mapped one-to-one to its envelope [3] and for the original real-valued data, the imaginary part of the analytic signal is uniquely computed via Hilbert transform. Therefore, the iterative adjustment of the spatial and temporal filter weights to obtain some specific envelope *e*_*m*_*t*) is theoretically equivalent to the adjustment of the weights to obtain this envelope’s generating analytic signal *s*_*m*_(*t*).

Assume that the training of the adaptive envelope detectors resulted in optimal spatial and temporal convolution weights 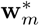 and 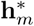 correspondingly. Now, perturb the spatial weights 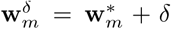 while keeping the temporal convolution weights vector at the optimum. Then, clearly, the optimal spatial weights can be found again as the solution to a convex optimization problem formulated over spatial subset of parameters:

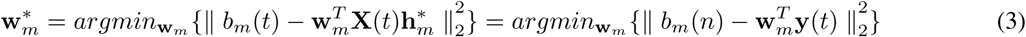

where the temporal weights are fixed at their optimal value, 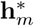 Similarly, when spatial weights are fixed at the optimal value 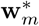, the temporal weights are expressed by the equation:

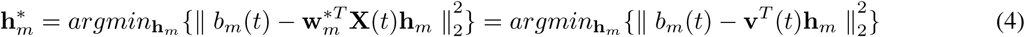

Given the forward model (1) and the regression problem (3) and assuming mutual statistical independence of the rhythmic potentials *s*_*m*_(*t*), *m* = 1, …, *M*, the topographies of the underlying neuronal populations can be found as [7, 5]

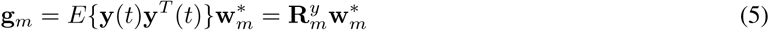

where **y**(*t*) = **X**(*t*)**h**_*m*_ is a temporally filtered vector of multichannel data and 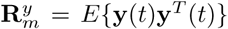 is a *K*× *K* spatial covariance matrix of the temporally filtered data, assuming that *x*_*k*_(*t*), *k* = 1, …, *K* are all zero-mean processes. Thus, when interpreting individual spatial weights corresponding to each of the *M* branches of the architecture shown in Figure 2 one has to take into account the temporal filter weights **h**_*m*_ this *m* th branch is tuned for. Therefore, to transform the spatial weights of different branches into spatial patterns, branch-specific spatial covariance matrices 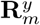 should be used that depend on the temporal convolution weights in each particular branch. This becomes obvious if one remembers that the spatial and temporal filtering operations are linear and can be interchanged. However, since each of the branches has its own temporal filter, it has to be situated past the spatial unmixing step which makes the obtained result less intuitive.

The temporal weights can be interpreted in a similar way. The temporal pattern is calculated as

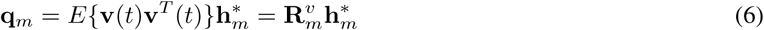

where 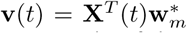 is a spatially filtered chunk of incoming data and 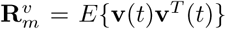is an *N* × *N* tap covariance matrix of the spatially filtered data, assuming that *x*_*k*_(*t*), *k* = 1, …, *K* are all zero-mean processes. As with the spatial patterns, when interpreting individual temporal weights corresponding to each of the *M* branches of the architecture shown in Figure 2, one has to take into account the spatial filter weights **w**_*m*_ used to feed the individual *m*−th branch. To transform the temporal convolution weights of different branches into temporal patterns, branch-specific tap covariance matrices 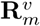 should be used that depend on the spatial point-wise convolution weights of each particular branch. To make sense out of the temporal pattern, we usually explore it in the frequency domain, i.e. 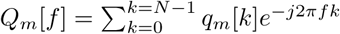 where *q*_*m*_ [*k*] if the *k*-the element of **q**_*m*_.

In the above for illustrative purposes we assumed that data chunk length *N* is equal to the length of the filters in the convolutional layers. In general, this does not have to be the case. Our assumption emphasizes the formal similarity between the spatial and temporal dimensions. Additionally, we emphasize that the approach to the interpretation of temporal patterns requires taking into account the correlation structure of the independent variable in the regression model. Relaxing the assumption about the filter length, equation (2) has to be rewritten using the convolution operation between the spatially filtered data and temporal convolutions weights **h**_*m*_. In this case, instead of a scalar, the equation returns a vector of samples, with the vector length depending on the choice of strategy used to deal with the transient at the edges of the chunk.

It may be easier to operate in the frequency domain from the very beginning, and use the standard Wiener filtering arguments. In the frequency domain, the Wiener filter weights can be expressed as a function of the power spectral density 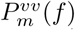 of the spatially filtered sensor data 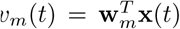 in the *m*-th branch and the density of cross-spectrum, 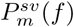, between *s*_*m*_(*t*) and *v*_*m*_(*t*):

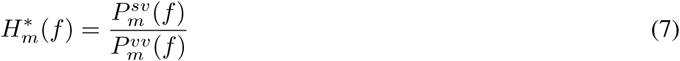

Then, using the assumption that *η*(*t*) and **s**(*t*) in (1) are statistically independent, we obtain the following expressions:

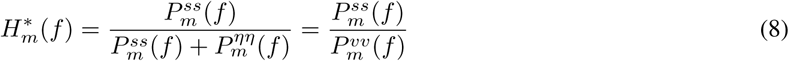

Therefore, the frequency-domain pattern of the signal isolated by the *m*-th branch spatial filter can be computed as

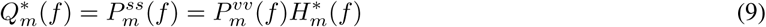

where 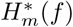 in (9) is the Fourier transform of the vector 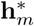 containing temporal-convolution weights identified during the adaptation of the envelope detector in the *m* − *th* branch. Viewing this result as a product of learning, we can say that learning to decode we gain access to the spectral pattern of activity (9) of the neuronal population critical for the particular decoding task. Importantly, the temporal filter weights alone will be informative of the population activity spectral pattern only in case the input timeseries are white.

The spatial patterns **g**_*m*_ of neuronal sources recovered from the spatial filtering weights [5] are routinely used for dipole fitting to localize functionally important neural sources [15]. The spectral patterns interpreted according to (9) and (6) can be used to fit the models of neural population dynamics, which are relevant to specific decoding tasks, see for example [16].

### 2.3 Simulations

To explore the performance of the proposed approach, we performed a set of simulations. The simulated data corresponded to the setting shown in the phenomenological diagram (Figure 1). We simulated *I* = 4 task related sources with rhythmic potentials *s*_*i*_(*t*). The potentials of these four task related populations were generated as narrow-band processes (in 30-80 Hz], 80-120 Hz], 120-170 Hz and 170-220 Hz bands) resulting from filtering Gaussian pseudo-random sequences with a bank of FIR filters. We then simulated the kinematics *z*(*t*), as a linear combination of the four envelopes of these rhythmic signals with randomly generated vector of coefficients.

We used *J* = 40 task-unrelated rhythmic sources with activation timeseries obtained similarly to the task-related sources but with filtering within the following four bands: 40-70 Hz, 90-110 Hz, 130-160 Hz, and 180 - 210 Hz bands. As a result, we obtained 10 task unrelated sources active in each of these bands making it total of *J* = 40 task unrelated sources. To simulate volume conduction effect, we randomly generated a 4×5 dimensional forward matrix **G** and a 40 ×5 dimensional forward matrix **A**. There matrices mapped the task-related and task-unrelated activity, respectively, onto the sensor space.

We generated 20 minutes worth of data sampled at 1,000 Hz and split them into two equal contiguous parts. We used the first part for training and the second for testing.

## 3 Experimental datasets

First, in order to compare the compact CNN architecture with the top linear models that rely on preset features, we used publicly available data collected by Kubanek et al. from the BCI Competition IV. This dataset contains concurrent multichannel ECOG and finger flexion kinematics measurements collected in three epileptic patients implanted with ECoG electrodes for medical reasons. The database consists of 400 seconds of training data and 200 seconds of test data. The recordings were conducted with 64 or 48 electrodes placed over the sensorimotor cortex. The exact spatial locations and the order of the electrodes were not provided. As a baseline in this comparison, we chose the winning solution offered by Nanying Liang and Laurent Bougrain [11]. This solution employs extracting the amplitudes of the data filtered in 1-60 Hz, 60-100 Hz, and 100-200 Hz band followed by a pairwise feature selection and decoded using Wiener filter with N = 25 taps from the immediate past.

The other dataset comes from our laboratory. The recordings were conducted with a 64-channel Adtech microgrid connected to EB Neuro BE Plus LTM Bioelectric Signals Amplifier System that sampled data at 2048 Hz. The amplifier software streamed data via Lab Streaming Layer protocol. The experimental software supported this protocol, implemented the experimental paradigm (a finger movement task) and synced ECoG and kinematics. Finger kinematics was captured by Perception Neuron system as relative angles for the sensor units attached to finger phalanges, and sampled at 120 Hz. Finger flexion-extension angle was used as kinematics timeseries, *z*(*t*).

The recordings were obtained in two patients with pharmaco-resistant form of epilepsy; ECoG electrodes were implanted for the purpose of pre-surgical localization of epileptic foci and mapping of eloquent cortex. Thus,for these data, unlike in the case of Berlin BCI IV competition data we knew cortical location of each electrode and could visualize spatial patterns of activity. The patients performed self-paced flexions of each individual finger for 1 min. The study was conducted according to the ethical standards of the 1964 Declaration of Helsinki. All participants provided written informed consent prior to the experiments. The ethics research committee of the National Research University, The Higher School of Economics approved the experimental protocol of this study

## 4 Results for simulated data

### 4.1 Adaptive envelop detector

As described in Methods, to interpret optimal temporal convolution weights we need to consider the spectral characteristics of neural recordings. To illustrate this, we trained a single-channel adaptive envelope detector in the environment with the interference occupying a subrange of the target signal frequency band. As can be seen from Figure 3, the Fourier profile of the identified temporal convolution weights can not be used to assess the power spectral density of the underlying signal as it has a characteristic deep over the frequency range occupied by the interference. The presence of this deep can be understood from the expression of Wiener filter transfer function (8). It shows that the transfer function will have a smaller then 1 gain over the frequency range where an interference is present, i.e: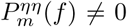. For simulations shown in Figure 3 the interference occupied 50-100 Hz frequency range. At the same time, the expression in (9) allows us to obtain a proper pattern that matches well the simulated spectral profile. Conversely, using the FFT of the convolutional filter weights yields fundamentally erroneous estimates of the frequency-domain patterns and potentially erroneous interpretation of the underlying neurophysiology.

**Figure 3:**
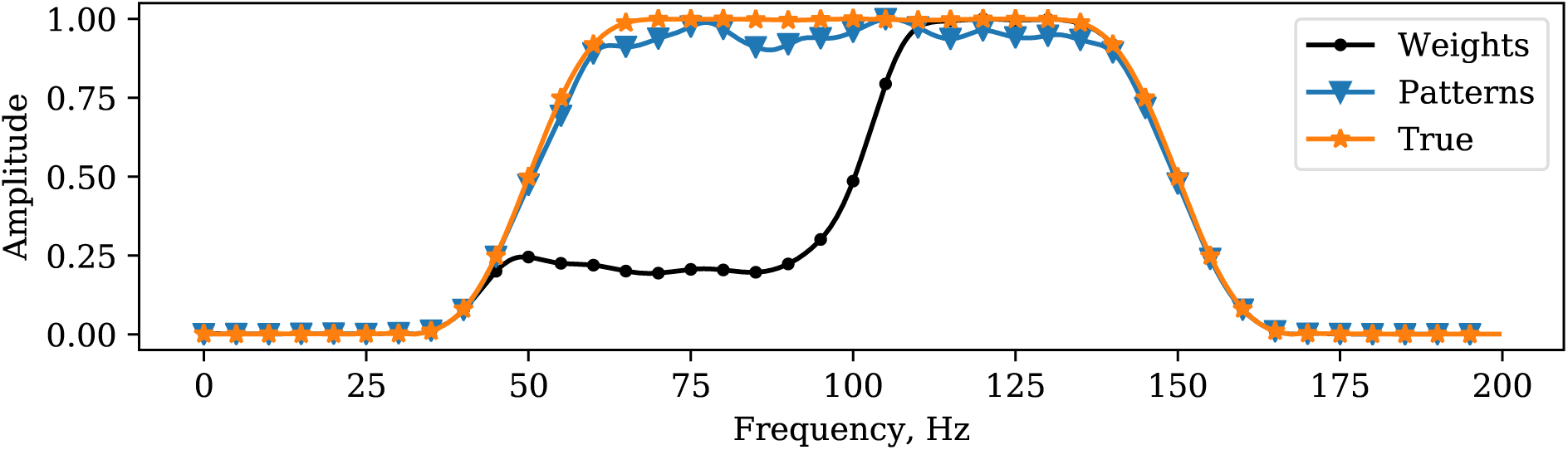
The importance of taking into account the power spectral density of the independent variable when interpreting linear regression weights. The true pattern (*) gets erroneously reproduced as the FFT of the temporal convolution weights (•). Taking into account the power spectral density of the spatially filtered signal allows to fix this situation (▾)

### 4.2 Realistic simulations

For the simulated data, we trained the algorithm to predict the kinematic variable *z*(*t*). In the noiseless case, the proposed architecture achieved accuracy of 99% measured as correlation coefficient between the true and the recovered kinematics, see Figure 4. We then compared the envelopes at each of the four branches of our architecture and observed that the true latent variable timeseries (in the form of the underlying narrow-band envelopes) matched very well those estimated with our architecture. Figure 5 shows estimated envelopes *r*_*m*_(*t*), superimposed on the true envelopes and the underlying narrow-band process *b*_*m*_(*t*) (Figure 2). We can see a good agreement between the estimated and true envelope timeseries with pairwise correlation coefficients within 87 - 96 % range.

**Figure 4:**
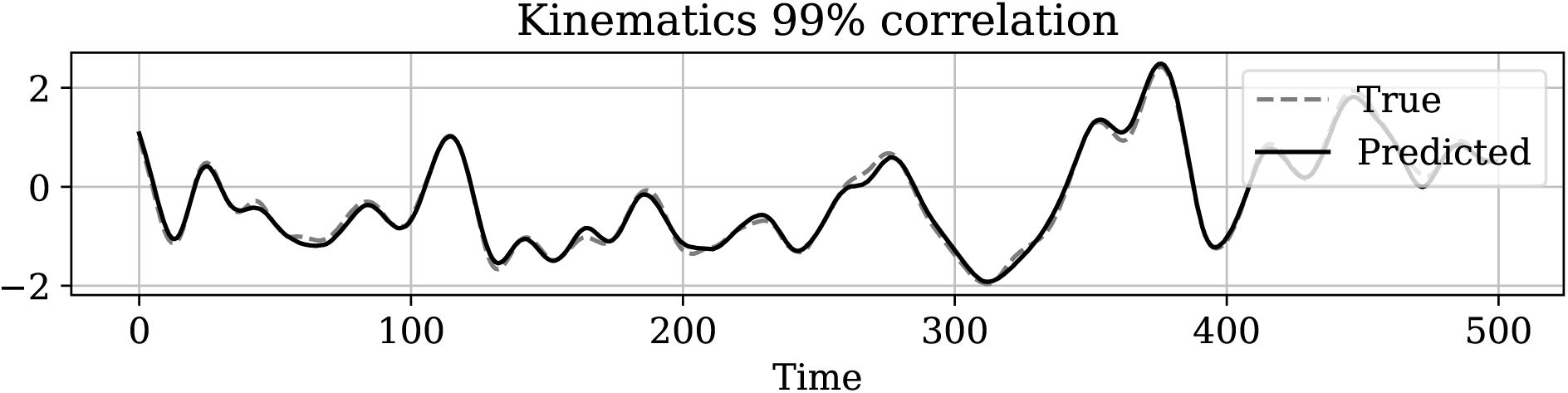
Realistic simulations. Actual and decoded kinematics.

**Figure 5:**
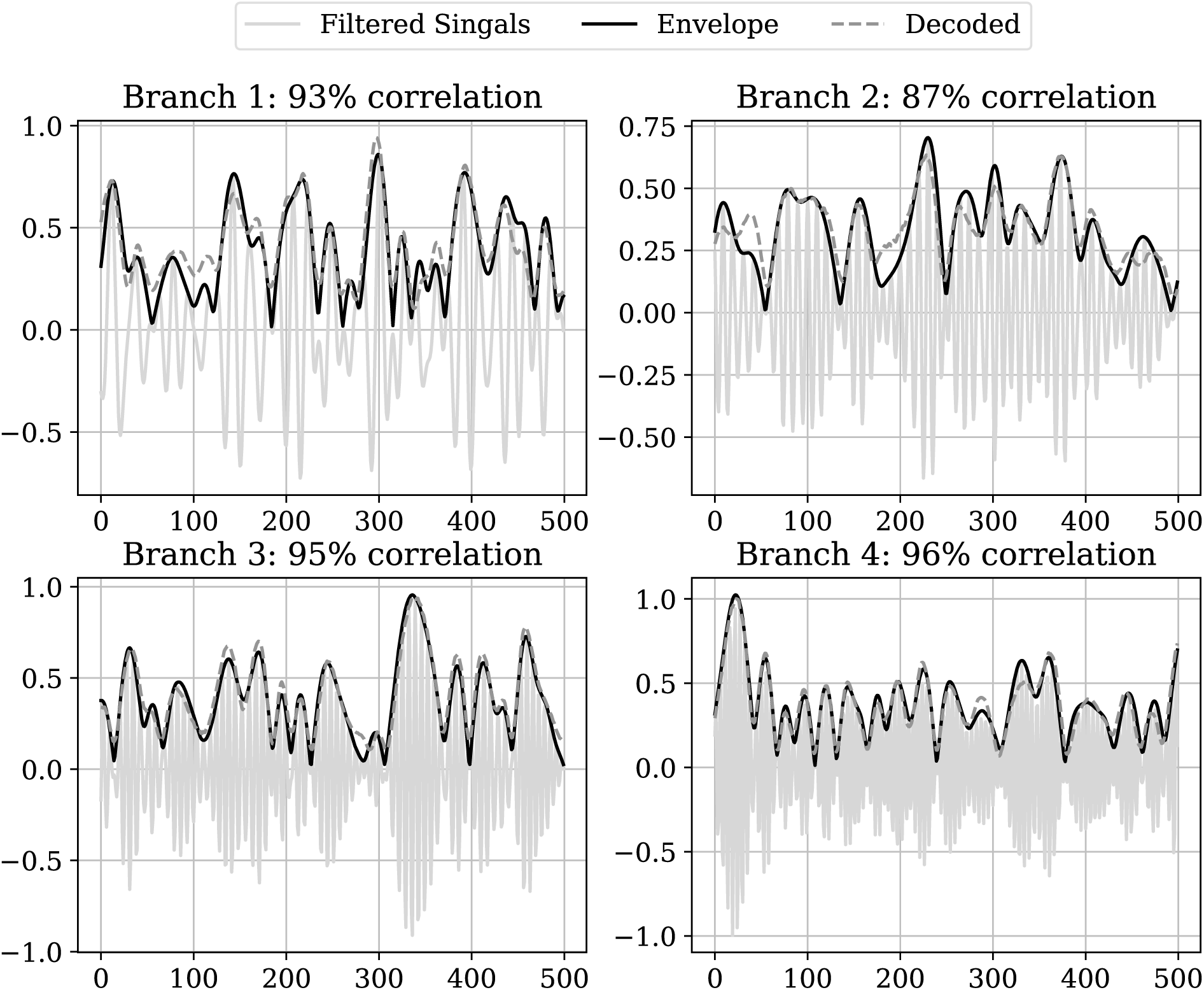
Branch envelopes decoding. Comparison between the true and the decoded envelope in each of the four branches.

As described in Methods, for spatial weights interpretation, we used the linear estimation theoretic approach [7, 5] To warn against its naive implementation in the context of architectures that combine spatial and temporal filtering, we computed spatial patterns where we used the input data covariance, **R**^*x*^, without taking into account the individual-branch temporal filters. In the corresponding plots, we refer to the patterns determined using this approach as *Patterns naive*. The proper way to apply this estimation approach is to compute spatial covariance, **R**^*y*^, for the temporally filtered data (6). These properly determined patterns are labeled as *Patterns* in the subsequent plots.

In the right column of Figure 6, we show the results of reconstructing spatial patterns in the noiseless case for all four branches of the network. As expected, the spatial *Patterns naive* and *Patterns* are identical and match the ground truth exactly. The left column shows Fourier representations of the temporal patterns and weights where we can observe that in the noise-free scenario Fourier representations of the temporal weights matches exactly the power spectral density of the simulated data.

**Figure 6:**
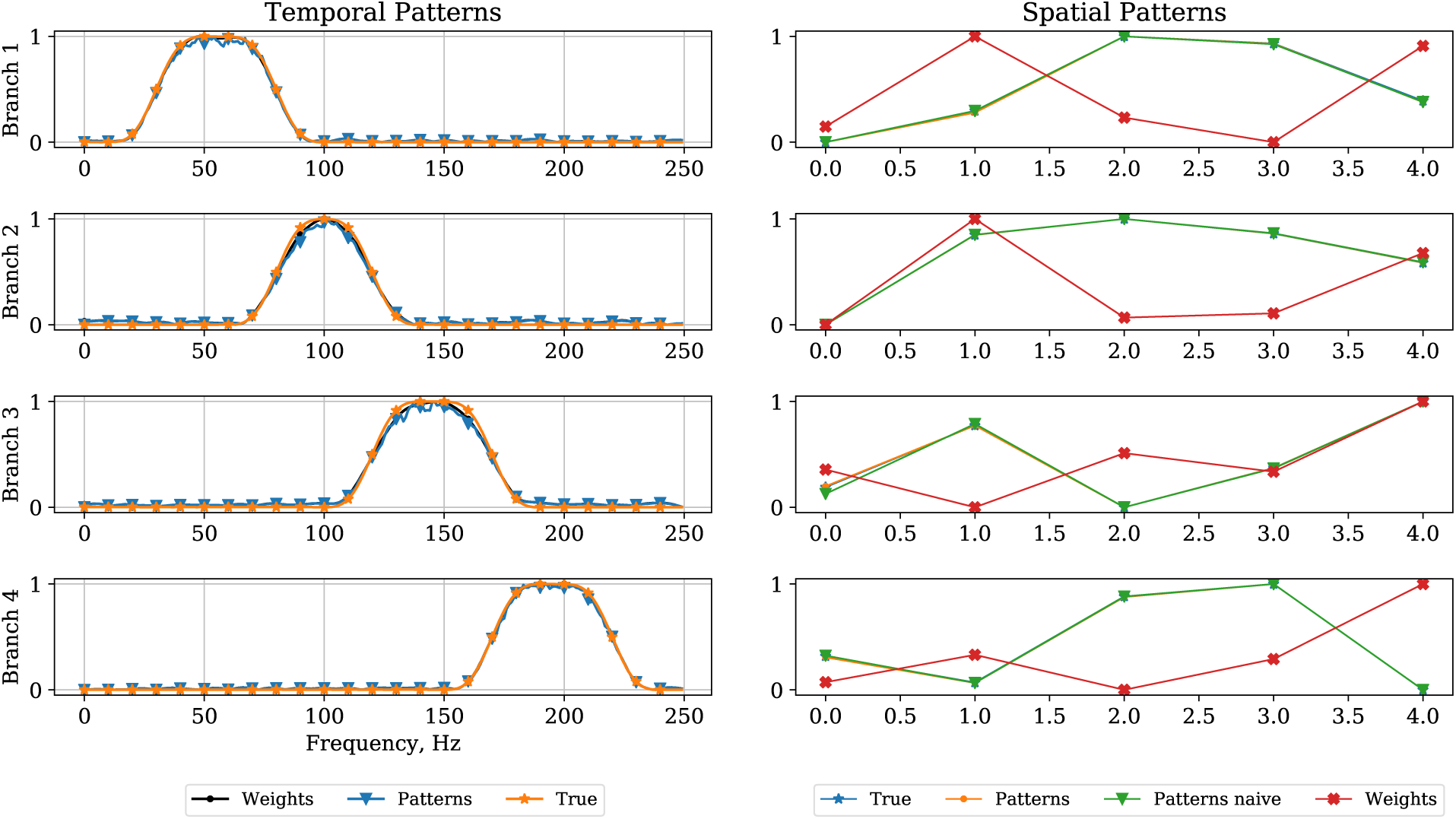
Temporal (left) and spatial(right) patterns obtained for the noiseless case. See the main text for description.

In the noisy case demonstrated in Figure 7, only *Patterns* match well with the simulated topographies of the underlying sources. Spectral characteristics of the trained temporal filtering weights exhibit characteristic deeps in the bands corresponding to the activity of the interfering sources. After applying expression (9), we obtain the spectral patterns that more closely match the simulated ones and have the deeps compensated.

**Figure 7:**
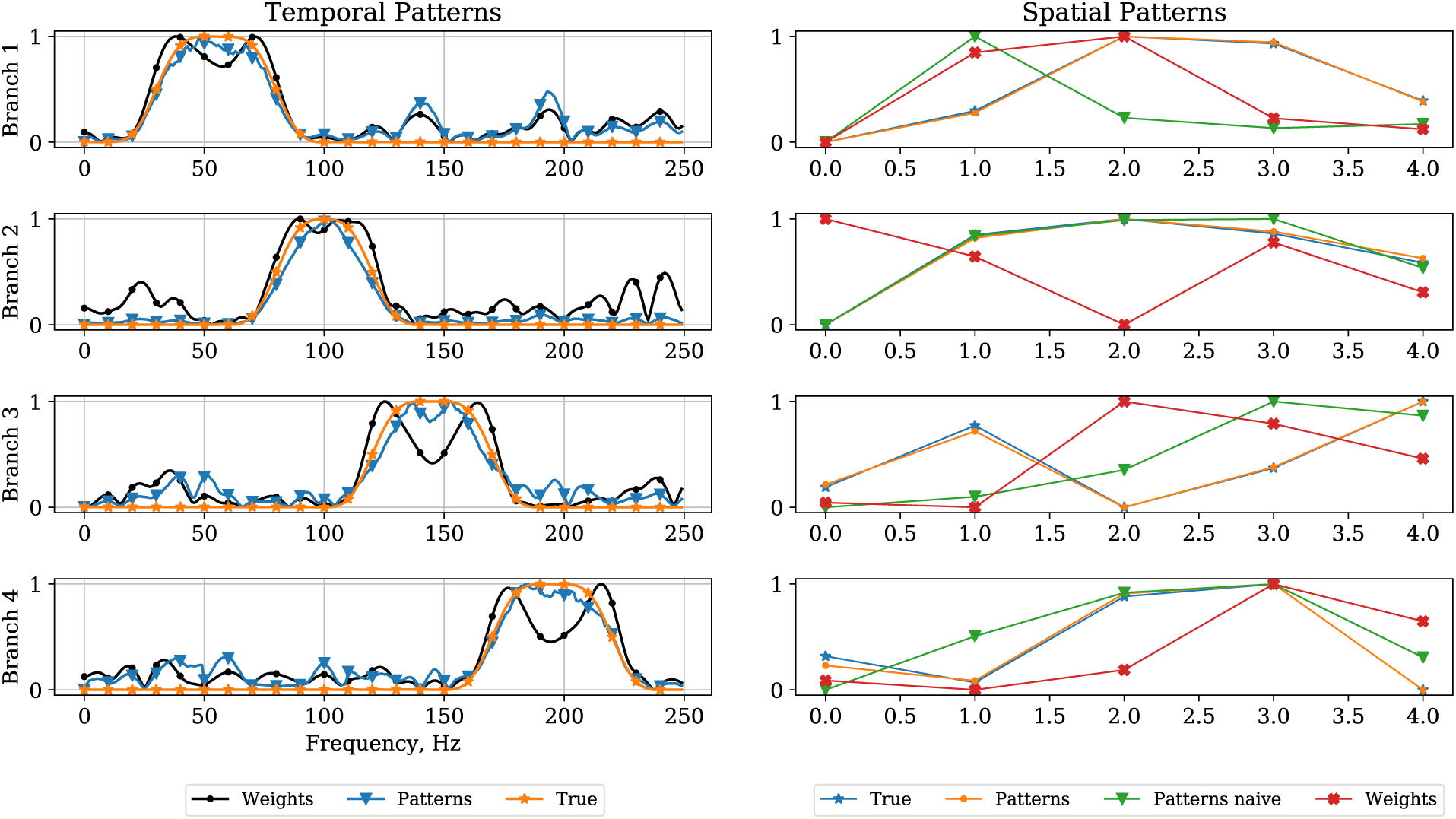
Temporal (left) and spatial(right) patterns obtained for the noisy case, SNR = 1.5. See the main text for description.

### 4.3 Monte-Carlo simulations

In the above plots we showed two specific cases of this architecture operating in the noisy and noiseless cases for a fixed spatial configuration of task related and task unrelated sources as modelled by matrices **G** and **A**. To test the proposed approach for weights interpretation we performed Monte-Carlo study with different spatial configuration of sources at each trial and for the four different noise levels. To implement this, matrices **G** and **A** which model the volume conduction effects at each Monte Carlo trial were randomly generated according to 𝒩 (0, 1) distribution. We created 20 minutes worth of data sampled at 1000 Hz. For neural network training we use Adam optimiser. We made about 15k steps. At 5k and 10k step we halved the learning rate to get more accurate patterns. In total, we have performed more then 3k simulations. For each realisation of the simulated data we have trained the algorithm to predict the kinematic variable *z*(*t*) and then computed the patterns of sources the individual branches of our architecture got “connected” to as a result of training. Figure 8 shows that only the spatial *Patterns* interpreted using branch-specific temporal filters (blue dots) match well the simulated topographies of the true underlying sources. The spectral patterns recovered using the proposed approach also appear to match well with the true spectral profiles of the underlying sources. Directly considering the Fourier coefficients of the temporal convolution weights results into generally erroneous spectral profiles (red triangles). For spatial patterns we also show the results for naively estimated patterns without taking into account branch-specific temporal filtering (green dots). Thus, using the proper spectral patterns of the underlying neuronal population it is now possible to fit biologically plausible models, e.g. [16], and recover true neurophysiologial mechanisms underlying the decoded process.

**Figure 8:**
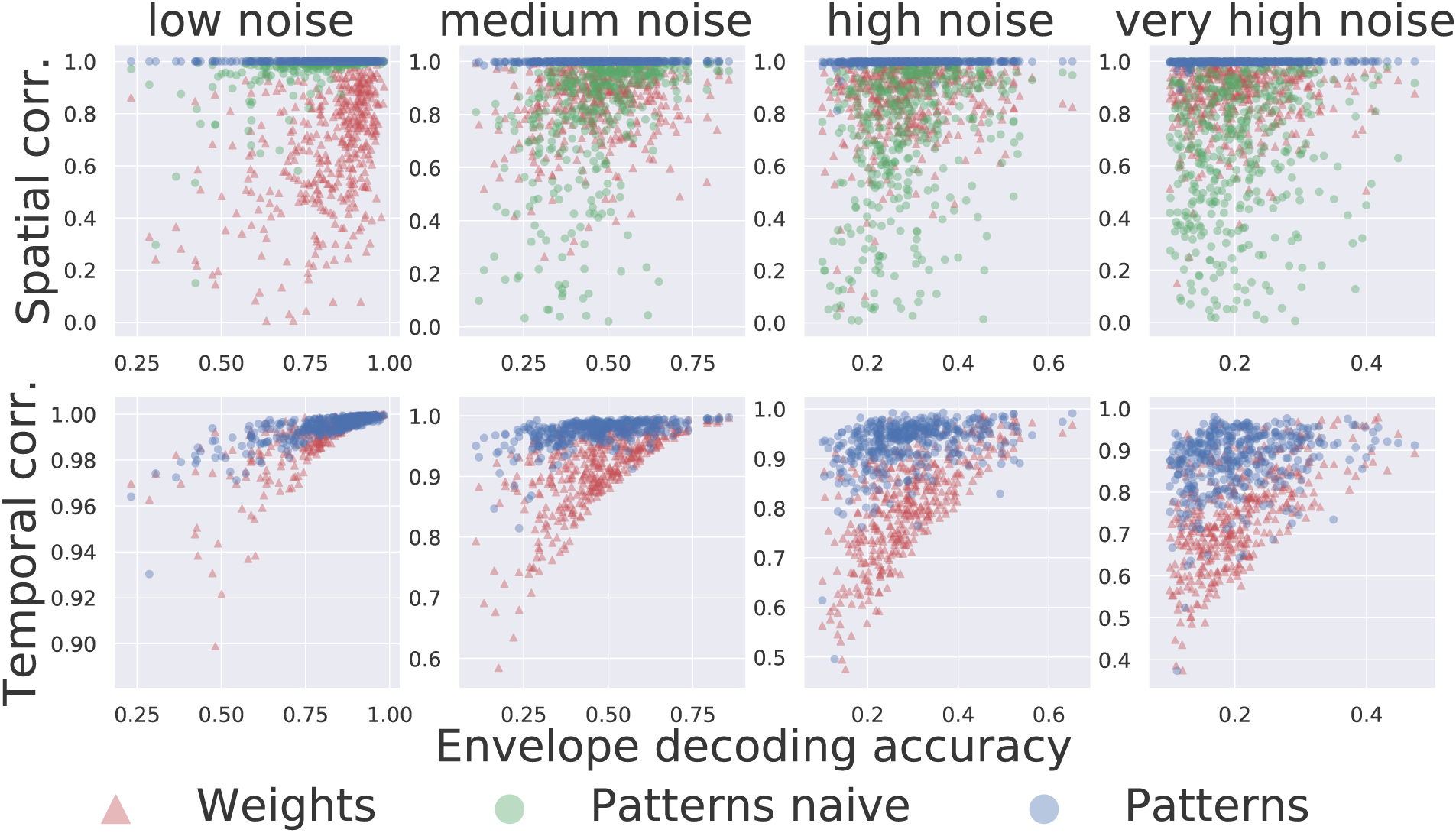
Monte-arlo simulations. Point coordinates reflect the achieved at each Monte Carlo trial envelope decoding accuracy (x-axis) and correlation coefficient with the true pattern (y-axis). Each point of a specific color corresponds to a single Monte Carlo trial and codes for a method used to compute patterns. *Weights* - direct weights interpretation, *Patterns naive* - spatial patterns interpretation without taking branch specific temporal filters into account, *Patterns* - the proposed method

## 5 Analysis of real experimental data

### 5.1 Berlin BCI Competition IV data

In the context of electrophysiological data processing, the major advantage of the architectures inspired by the deep-learning principle is their ability to automatically select features while performing classification or regression tasks [21]. When applied to to the data from Berlin BCI Competition IV, the compact CNN architecture – based on the adaptive envelope detectors –performed on par or better than the winning solution by Lian and Bougrain [11], see Table 1.

**Table 1:** Comparison of the performance of the proposed architecture (NET) and the winning solution (Winner) of Berlin BCI IV competition dataset(4)

| Subject 1 |  |  |  |  |  |
| --- | --- | --- | --- | --- | --- |
|  | Thumb | Index | Middle | Ring | Little |
| Winner | 0,58 | 0,71 | 0,14 | 0,53 | 0,29 |
| NET | 0,54 | 0,7 | 0,2 | 0,58 | 0,25 |

| Subject 2 |  |  |  |  |  |
| --- | --- | --- | --- | --- | --- |
|  | Thumb | Index | Middle | Ring | Little |
| Winner | 0,51 | 0,37 | 0,24 | 0,47 | 0,35 |
| NET | 0,5 | 0,36 | 0,22 | 0,4 | 0,23 |

| Subject 3 |  |  |  |  |  |
| --- | --- | --- | --- | --- | --- |
|  | Thumb | Index | Middle | Ring | Little |
| Winner | 0,59 | 0,51 | 0,32 | 0,53 | 0,2 |
| NET | 0,71 | 0,48 | 0,5 | 0,52 | 0,61 |

### 5.2 Dataset 2

We also applied the proposed solutions to the recordings that we conducted in two patients implanted with 8×8 ECoG microgrids grids placed over the sensorimotor cortex.

The following table shows the accuracy achieved with the proposed architecture for the decoding of finger movements. Figures 9 and 10 depict the interpretation of the obtained spatial and temporal weights. The plots are shown for the finger with the highest decoding accuracy (highlighted in bold in Table 2) for two patients.

**Table 2:** Decoding performance achieved in the two CBI patients. The table show correlation coefficient between the actual and the decoded finger trajectory for the four fingers in two patients.

|  | Thumb | Index | Ring | Little |
| --- | --- | --- | --- | --- |
| Subject 1 | 0,48 | **0,79** | 0,61 | 0,32 |
| Subject 2 | 0,73 | 0,55 | 0,78 | **0,79** |

**Figure 9:**
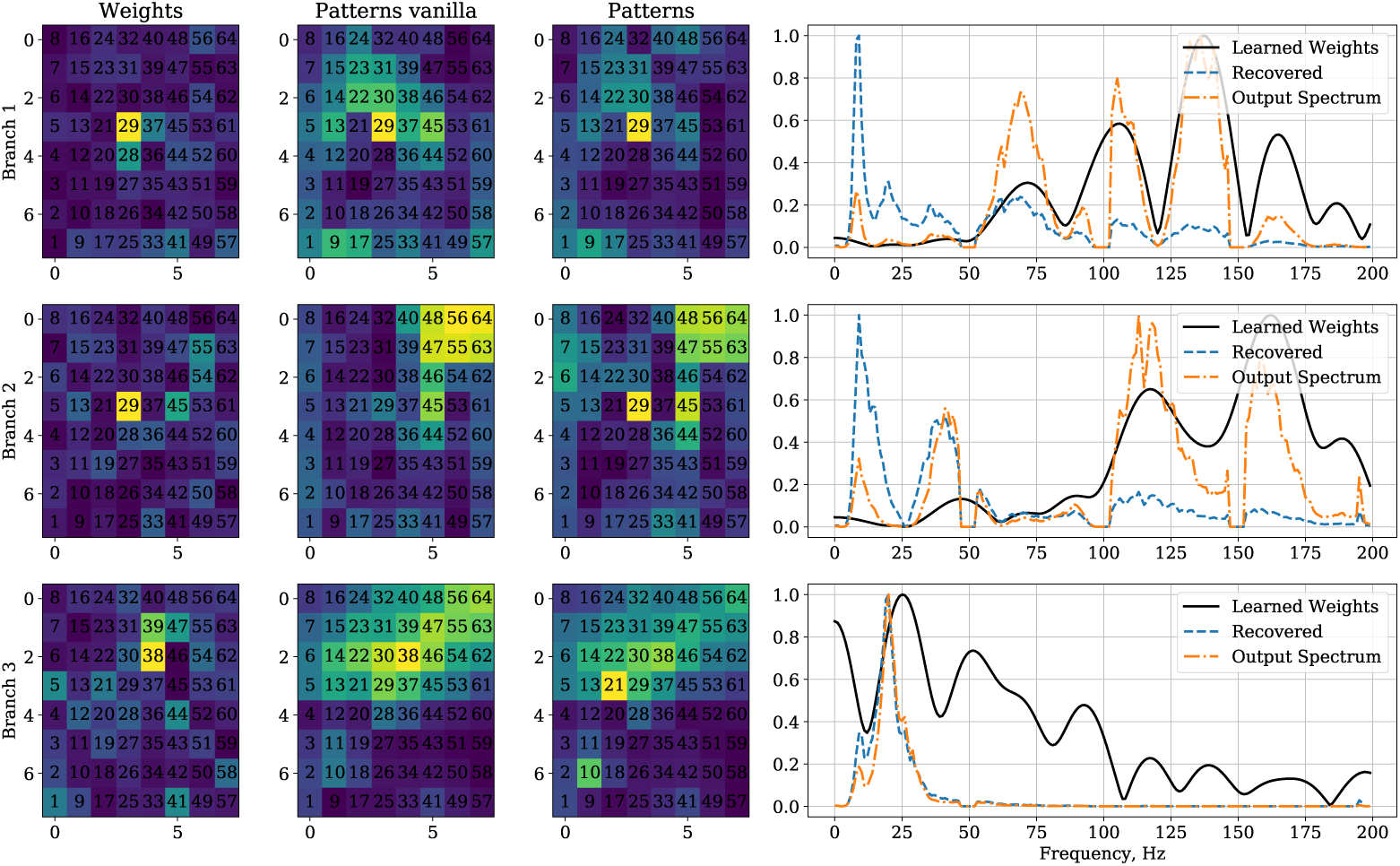
Network weights interpretation for the index finger decoder in CBI patient 1. Each row of plots corresponds to one of the three branches of the trained decoder. The left most column shows color-coded spatial filter weights, next two columns correspond to naively and properly recovered spatial patterns. The fourth column interprets temporal filter weights in the Fourier domain. Filter weights - solid line, power spectral density (PSD) pattern of the underlying LFP - dashed line. Another dashed line more similar to the filter weights Fourier coefficients is the PSD of the signal at the output of the temporal convolution block.

**Figure 10:**
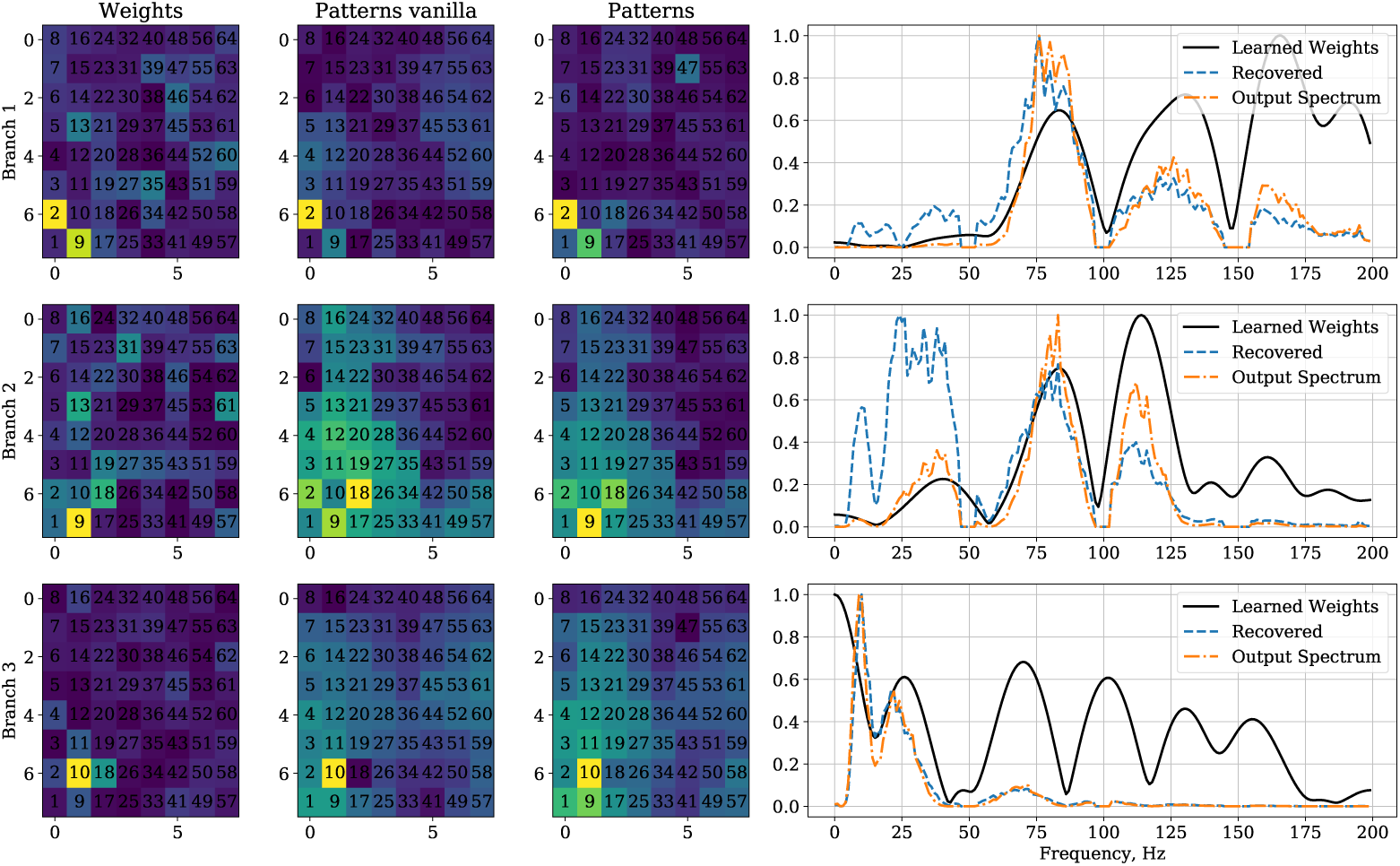
Network weights interpretation for the little finger decoder in CBI patient 2. Each row of plots corresponds to one of the three branches of the trained decoder. The left most column shows color-coded spatial filter weights, next two columns correspond to naively and properly rcovered spatial patterns. The fourth column interprets temporal filter weights in the Fourier domain. Filter weights - solid line, power spectral density (PSD) pattern of the underlying LFP - dashed line. Another dashed line more similar to the filter weights Fourier coefficients is the PSD of the signal at the output of the temporal convolution block.

The decoding architecture for both patients had three branches and each branch was tuned to a source with specific spatial and temporal patterns. In Figure 9, we show the spatial filter weights, naive patterns and proper patterns interpreted using the expression described in the the Methods section. It can be seen that, while the temporal filter weights (solid line) clearly emphasized the frequency range above 100 Hz in the first two branches, the actual spectral pattern of the source (dashed line) in addition to the gamma-band content had a peak at around 11 Hz (1st, 2^nd^ branches) and in the 25-50 Hz range (2nd branch). These peaks likely correspond to the sensorimotor rhythm and low-frequency gamma rhythms, respectively.The third branch appears to capture the lower-frequency range and its spatial pattern is noticeably more diffuse than that in the first two branches that capture the higher-frequency components. Similar observations can be made from Figure 10 that shows to the decoding results for the little finger in patient 2. Interestingly, the second branch frequency domain pattern (dashed blue line) appears to be significantly different from that obtained by a simple FFT of the weights vector (solid black line) and contains contributions from the lower 20-45 Hz frequency range. When fitting dynamical models of population activity to this recovered frequency domain pattern, the low-frequency components are likely to significantly affect the parameters of the corresponding dynamical model.

## 6 Conclusion

We first described a neurophysiologically interpretable architecture based on a compact convolutional network similar to those reported earlier [8, 23, 28]. Using this architecture, we extended the weights interpretation approach [5] to the interpretation of the temporal convolution weights and adapted it to the architectures with multiple branches with specific spatial and temporal filters in each. We tested the proposed approach using realistically simulated and experimental data. In the realistically simulated data, our architecture recovered with high accuracy the neuronal substrate that contributed to the simulated kinematics data. Interestingly, as shown in Figure 8, even in cases when the decoding accuracy was low, the spatial and temporal patterns appeared to be accurately recovered.

At the same time, the accuracy of spatial patterns recovery was noticeably higher than that for the temporal patterns (interpreted in the frequency domain). Such a behavior may partly stem from the fact that in our simulations the spatial patterns of each branch were encoded using only 5 coefficients as compared to 100-tap long temporal filters. One possible workaround that would reduce the number of the temporal convolution parameters to be identified is to use sinc-layer [19] applied in a network for analysis of acoustic signals. At the same time, this may reduce the flexibility in spectral shaping of individual branches of the network and require more pathways to be added to the network to achieve performance comparable to that reported here which may complicate the interpretation of the obtained decision rule.

We also applied the described architecture to Berlin BCI IV competition data. The compact CNN-based architecture delivered similar or better decoding accuracy as compared to the winning solution of this BCI competition [11]. In contrast to the traditional approaches, this architecture did not require any preset features. Instead, after the architecture was trained to decode finger kinematics, we could interpret its weights and extracted physiologically meaningful patterns corresponding to both spatial and temporal convolution weights. This latter was demonstrated using our own ECoG data collected from subjects performing a self-paced finger-flexion task for which we knew the spatial locations of electrodes.

## 7 Acknowledgment

This work is supported by the Center for Bioelectric Interfaces NRU HSE, RF Government grant, ag. No.14.641.31.0003.

